# Human Embryonic Stem Cell-derived Lung Organoids: a Model for SARS-CoV-2 Infection and Drug Test

**DOI:** 10.1101/2020.08.10.244350

**Authors:** Rongjuan Pei, Jianqi Feng, Yecheng Zhang, Hao Sun, Lian Li, Xuejie Yang, Jiangping He, Shuqi Xiao, Jin Xiong, Ying Lin, Kun Wen, Hongwei Zhou, Jiekai Chen, Zhili Rong, Xinwen Chen

## Abstract

The coronavirus disease 2019 (COVID-19) pandemic is caused by infection with the severe acute respiratory syndrome coronavirus 2 (SARS-CoV-2), which is spread primary via respiratory droplets and infects the lungs. Currently widely used cell lines and animals are unable to accurately mimic human physiological conditions because of the abnormal status of cell lines (transformed or cancer cells) and species differences between animals and humans. Organoids are stem cell-derived self-organized three-dimensional culture *in vitro* and model the physiological conditions of natural organs. Here we demonstrated that SARS-CoV-2 infected and extensively replicated in human embryonic stem cells (hESCs)-derived lung organoids, including airway and alveolar organoids. Ciliated cells, alveolar type 2 (AT2) cells and rare club cells were virus target cells. Electron microscopy captured typical replication, assembly and release ultrastructures and revealed the presence of viruses within lamellar bodies in AT2 cells. Virus infection induced more severe cell death in alveolar organoids than in airway organoids. Additionally, RNA-seq revealed early cell response to SARS-CoV-2 infection and an unexpected downregulation of ACE2 mRNA. Further, compared to the transmembrane protease, serine 2 (TMPRSS2) inhibitor camostat, the nucleotide analog prodrug Remdesivir potently inhibited SARS-CoV-2 replication in lung organoids. Therefore, human lung organoids can serve as a pathophysiological model for SARS-CoV-2 infection and drug discovery.

## Introduction

The current fast-evolving coronavirus disease 2019 (COVID-19) pandemic is caused by the severe acute respiratory syndrome coronavirus 2 (SARS-CoV-2), which infects lungs and can lead to severe lung injury, multiorgan failure, and death^1-3^. To prevent and effectively manage COVID-19, public health, clinical interventions, basic research, and clinical investigation are all emergently required. For basic research, it is essential to establish models that can faithfully reproduce the viral life cycle and mimic the pathology of COVID-19.

Cell lines and animals are two major models for coronavirus infection *in vitro* and *in vivo*, respectively^4-7^. Cell lines can be used to amplify and isolate viruses (like Vero and Vero E6 cells^8,9^), to investigate the viral infection (like primary human airway epithelial cells, Caco-2 and Calu-3 cells^3,5,10,11^), and to evaluate therapeutic molecules (like Huh7 and Vero E6 cells^12^). Animal models can be used to mimic tissue-specific and systemic virus-host interaction and reveal the complex pathophysiology of coronaviruses-induced diseases^7^. Mice, hamster, ferrets, cats, and non-human primates have been reported to model COVID-19^13-21^. These cell and animal models have greatly enriched our understanding of coronaviruses and assisted in the development of a variety of potential therapeutic drugs^7^. However, these models yet have obvious limitations. Species differences make animal model results unable to be effectively translated into clinical applications^22,23^. Species differences (cells from species other than humans, like Vero cells) and abnormal status (transformed or cancer cells) make cell models unable to faithfully reproduce the viral infection cycle and host response^24-26^.

Organoids are a three-dimensional structure formed by self-assembly of stem cells *in vitro*^27,28^. As the cell composition, tissue organization, physiological characteristics, and even functions are similar to natural organs in the body, organoids have been used for human virus studies^29,30^. For SARS-CoV-2, lung, kidney, liver, intestine, and blood vessel organoids have been reported to be sensitive for virus infection^31-37^. Here using human embryonic stem cells (hESCs)-derived lung airway and alveolar organoids, we demonstrate that SARS-CoV-2 infects ciliated cells, alveolar type 2 cells (AT2 cells) as well as rare club cells, and remdesivir is more potent than camosat to inhibit virus infection.

## Results

### Generation of human lung airway and alveolar organoids from hESCs

Based on our previous protocol^38^, as well as other reported protocols^39,40^, we developed an optimized method to differentiate human airway organoids (hAWOs) and alveolar organoids (hALOs) from hESCs, which contained six stages, embryonic stem cells (ESCs), definitive endoderm (DE), anterior foregut endoderm (AFE), ventralized anterior foregut endoderm (VAFE), lung progenitors (LPs), and hAWOs and hALOs (Fig. 1a, b). Quantitative RT-PCR revealed the expression dynamics of marker genes along differentiation (Fig. 1c). *POU5F1* (ESCs), *SOX17* (DE), *SOX2* (ESCs and lung proximal progenitors), *SOX9* (lung distal progenitors), *FOXA2* (lung epithelial cells), *NKX2*.*1* (lung epithelial cells), *P63* (basal cells), *SCGB1A1* (club cells), *MUC5AC* (goblet cells) and *SPC* (AT2 cells) showed expected expression patterns (Fig. 1c). Human lung organoids (hLOs) at day21 expressed lung and pan epithelial markers NKX2.1 and E-CAD, respectively (Fig. 1d). Immunofluorescent staining revealed that hAWOs contained basal cells (P63^+^), ciliated cells (acetylated TUBULIN, a-TUB^+^), club cells (CC10^+^), and goblet cells (MUC5AC^+^), as well as lung proximal progenitors (SOX2^+^) and proliferating cells (Ki67^+^) (Fig. 1e). And hALOs contained AT2 cells (SPC^+^) and AT1 cells (PDPN^+^ or AQP5^+^) (Fig. 1f). Since ACE2 is the receptor for SARS-CoV-2 for host cell entry and TMPRSS2 is the serine protease for spike (S) protein priming^5,9^, we checked their expression along the differentiation and found they were highly expressed in hAWOs and hALOs (Fig. 1g).

**Fig.1.**
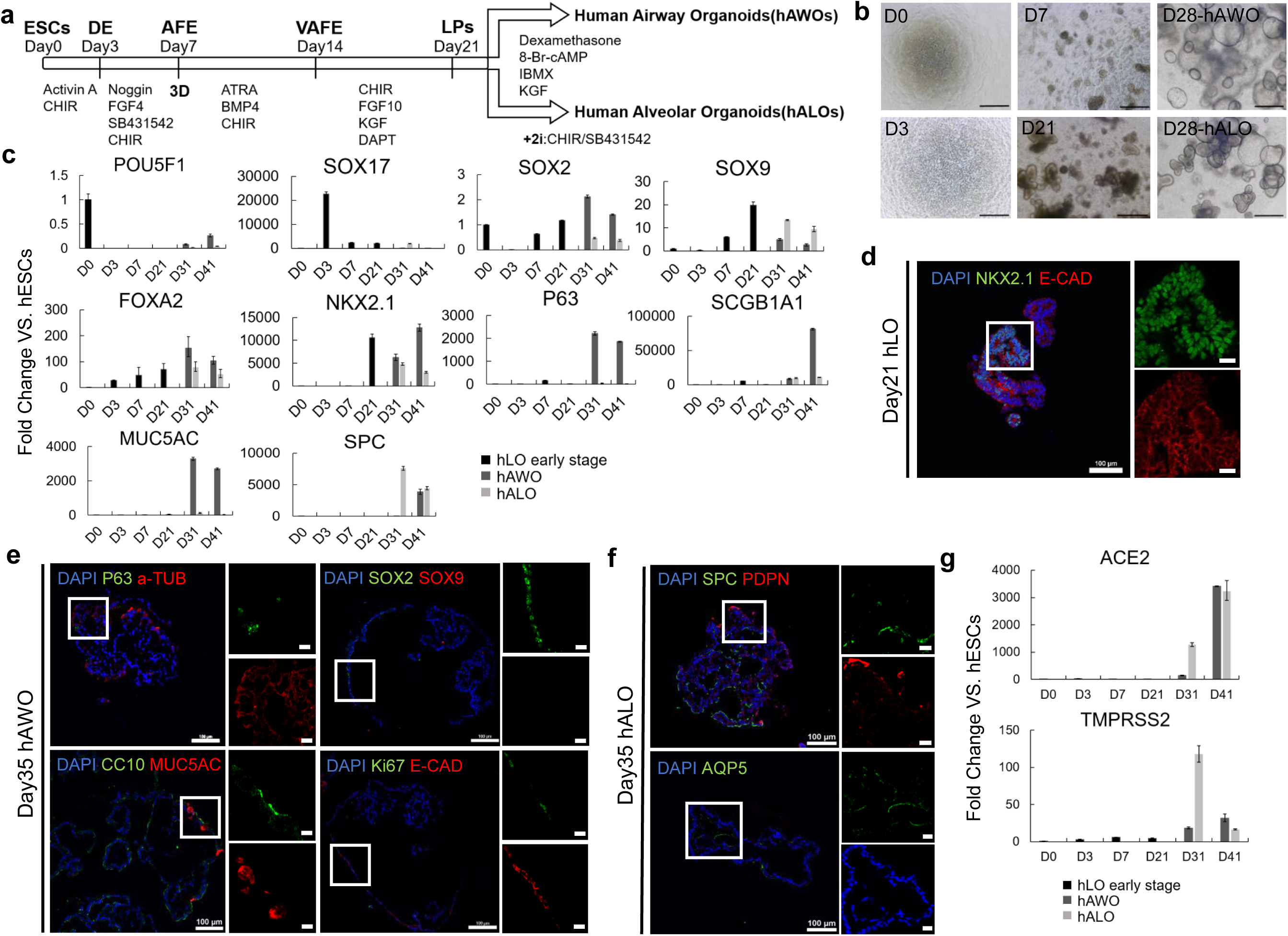
Generation of human airway and alveolar organoids from hESCs. **a**. Schematic of differentiation protocol and stages from hESCs to human airway organoids (hAWOs) and human alveolar organoids (hALOs). **b**, Representative images at the indicated differentiation stages. Scale bar, 500 μm. **c**, Fold change of lineage marker genes from day 0 (D0) to D41 over undifferentiated hESCs by quantitative RT-PCR (2^-ΔΔCt^). D0-D21, hLOs early stage. D21-D41, organoids split into two groups with different differentiated medium (hAWOs and hALOs). *POU5F1*, embryonic stem cell marker, *SOX17*, definitive endoderm marker, *SOX2*, embryonic stem cell and proximal airway cell marker, *SOX9*, distal alveolar progenitor cell marker, *FOXA2* and *NKX2*.*1*, lung progenitor lineage marker, *P63*, basal cell marker, *SCGB1A1 (CC10)*, club cell marker, *MUC5AC*, goblet cell marker, *SPC*, AT2 cell marker. Normalized to *GAPDH*. Bars represent mean ± SD, n=3. **d-f**, Cell lineage marker expression in human lung progenitor organoids (hLOs), human airway organoids (hAWOs), and human alveolar organoids (hALOs). Immunofluorescence images of NKX2.1 and E-Cadherin (epithelial cells) expression in D21 hLOs (**d**), of P63, SOX2, CC10, Ki67 (proliferation cells) and acetylated tubulin (ciliated cells), SOX9, MUC5AC, E-Cadherin protein expression in D35 hAWOs (**e**), and of SPC, AQP5 (AT1) and PDPN (AT1) expression in D35 hALOs (**f**). Nuclei were counterstained with DAPI. Scale bar, 100μm (left panel); 20μm (right panel). Boxes represent zoom views. **g**, Fold change of *ACE2* and *TMPRSS2* gene expression from D0 to D41 over undifferentiated hESCs by quantitative RT-PCR (2^-ΔΔCt^). Normalized to *GAPDH*. Bars represent mean ± SD, n=3.

### SARS-CoV-2 infects human airway and alveolar organoids

To test whether SARS-CoV-2 infects human lung organoids, hAWOs and hALOs (ranging from day 31 (D13) to D41) were exposed to SRAS-CoV-2 at a multiplicity of infection (MOI) of 1. Samples were harvested at indicated time points after infection and processed for the various analyses shown in Fig 2-5. Live virus titration on Vero E6 cells and quantitative RT-PCR of viral RNA in the culture supernatant and cell lysates showed that hAWOs and hALOs were productively infected by SARS-CoV-2 (Fig. 2a, b). Viral RNA and infectious virus particles could be detected as early as 24 hours post infection (hpi), increased at 48 hpi, and remained stable at 72 hpi. Compared to hALOs, hAWOs produced less virus at 24 hpi and similar amount of virus at 48 hpi and 72 hpi (Fig. 2a, b). Co-immunostaining of viral nucleocapsid protein (NP) and pan epithelial marker E-CAD showed that SARS-CoV-2 infected epithelial cells in human lung organoids (Fig. 2c). Quantification analysis showed that the percentages of infected hAWOs increased from about 50% at 24 hpi to about 75% at 72 hpi (Fig. 2d). And the percentages of infected cells within a single hAWO increased from about 24.9±3.7% at 24 hpi to 63.9±6.1% at 72 hpi (Fig. 2e). For hALOs, the percentages of infected organoids remained stable at about 85% and the percentages of infected cells per organoid remained about 30%-40% from 24 hpi to 72 hpi. These cellular infection results were consistent with viral RNA detection and infectious viral particle titration results.

**Fig.2.**
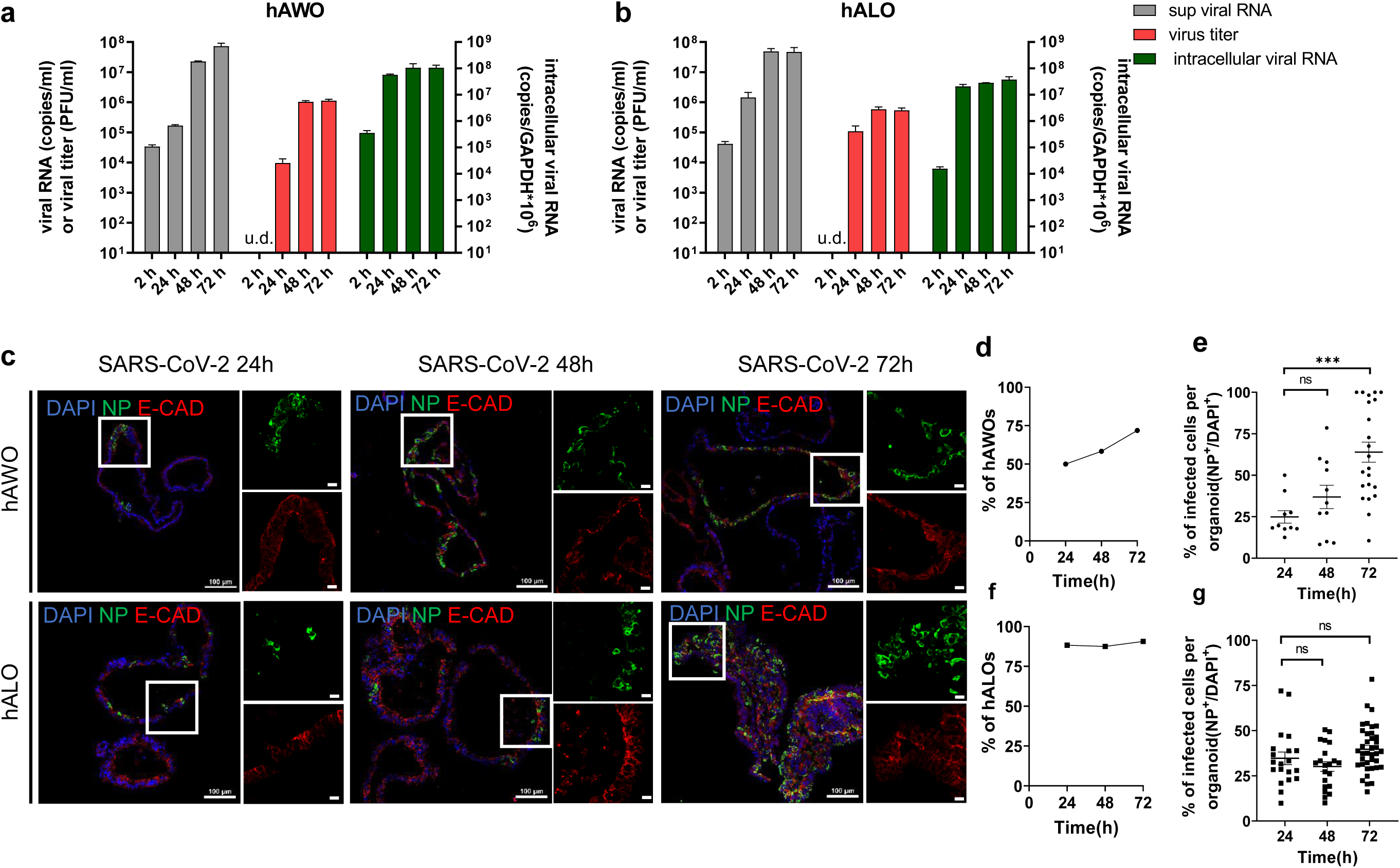
SARS-CoV-2 replicates in human airway and alveolar organoids. **a, b**, The viral RNA and virus titer in the culture supernatant and relative intracellular viral RNA in cell lysates in hAWOs (**a**) and hALOs (**b**) were detected at indicated time points post infection. **c**, Immunofluorescence images of viral nucleoprotein (green) and epithelial marker E-cadherin (red) expression with DNA stain (DAPI, blue) in SARS-CoV-2 infected hAWOs and hALOs. Scale bar, 100μm (left panel); 20μm (right panel). Boxes represent zoom views. **d, f**, Percentage of hAWOs (**d**) and hALOs (**f**) harboring SARS-CoV2 infected cells at different time points. At least 30 different organoids were counted per condition. **e, g**, Percentage of infected cells per infected hAWOs (**e**) and hALOs (**g**). At least 10 organoids were counted in **e** and at least 20 organoids in **g**. \*\*\**p* <0.001, by one-way ANOVA analysis.

### SARS-CoV-2 infects ciliated cells, alveolar type 2 cells and rare club cells

To determine the cell tropism of SARS-CoV-2, we co-stained each cell lineage marker with viral N protein and virus receptor ACE2. Microscopy analyses revealed that ciliated cells (a-TUB^+^) and alveolar type 2 cells (Pro-SPC^+^) were the major target cells (Fig.3a, b, and Fig. S1), which was consistent with the previous report^41^. In addition, rare club cells (CC10^+^) could be infected (Fig.3a). In hAWOs, about 90%-95% infected cells were ciliated cells and about 5%-10% were club cells, and no basal (P63^+^) or goblet cells (MUC5AC^+^) were found infected (Fig. 3c). In hALOs, 100% infected cells were AT2 cells and no AT1 cells (PDPN^+^) were found infected (Fig. 3c). We also measured the percentages of infected cells within ciliated cells and AT2 cells. About 26±3.6% at 24 hpi and 64.5±9.8% at 72 hpi of ciliated cells were infected, and the percentages of infected AT2 cells remained stable at about 30%-40% from 24 hpi to 72 hpi (Fig. 3d, e). The distinct infection dynamics of ciliated cells and AT2 cells indicated that more and more ciliated cells could be infected by SARS-CoV-2 during a prolonged infection period and even all the ciliated cells could be finally infected when given long enough infection time. On the contrary, only a subpopulation of AT2 cells (about 30-40%) was sensitive for viral infection although they could be quickly infected (within 24 hpi). The identity of the SARS-CoV-2 sensitive AT2 cell subpopulation and why other AT2 cells could not be infected need further investigation.

**Fig.3.**
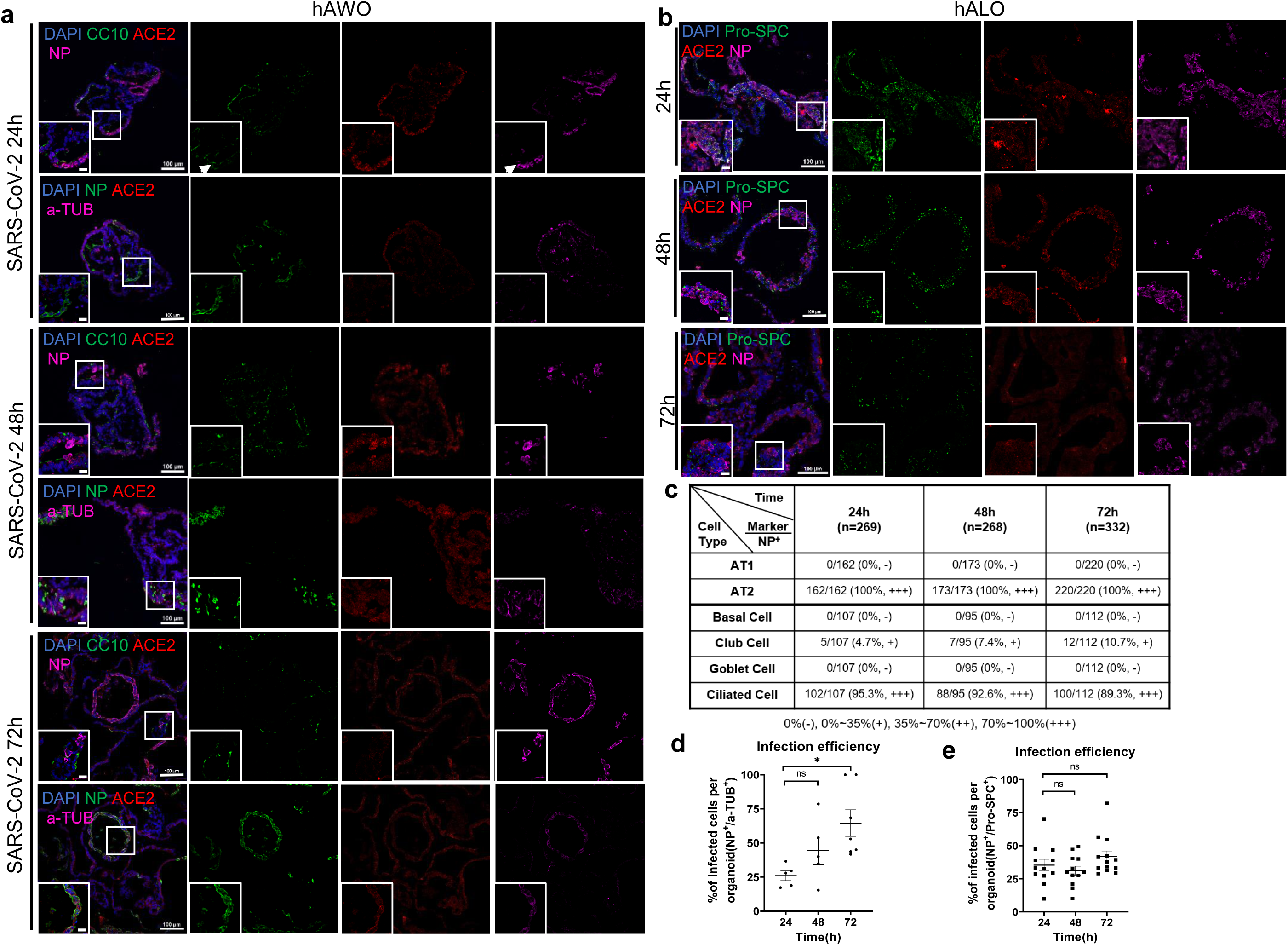
SARS-CoV-2 infects ciliated cells, alveolar type 2 cells and rare club cells. **a, b**, Representative immunofluorescence images of nucleoprotein, ACE2 and indicated cell linage marker expression with DNA stain (DAPI). Club cells (CC10^+^) and ciliated cells (acetylated Tubulin^+^) were stained in human airway organoids at indicated time points (**a**). Arrowheads indicate infected club cells. Alveolar type 2 cells (pro-SPC^+^) were stained in human alveolar organoids (**b**). Scale bars, 100µm; bottom left corner, 20µm. Boxes represent zoom views. **c**, Summary of the percentage of different cell types to SARS-CoV-2 infected cells in human airway and alveolar organoids. 269, 268, and 332 nucleoprotein positive cells were counted at 24, 48, and 72 hpi, respectively, after infection. -, negative for the nucleoprotein staining; +, 0%∼35% positive for the nucleoprotein; ++, 35%∼70% positive for the nucleoprotein; +++, 70%∼100% positive for the nucleoprotein. **d, e**, Percentage of infected ciliated cells (acetylated Tubulin^+^) per infected airway organoid (**d**) and infected alveolar type 2 cells (pro-SPC^+^) per alveolar organoid (**e**). At least 5 organoids were counted in **d** and at least 13 organoids in **e**. * *p* <0.05, by one-way ANOVA analysis.

We noted that viral infected cells expressed ACE2 but not all ACE2 expressing cells were infected. TMPRSS2 is another known factor that determines SARS-CoV-2 cell entry^5^, and therefore we checked the expression pattern of TMPRSS2 in human lung organoids. Immunostaining analyses showed that TMPRSS2 was ubiquitously expressed in both hAWOs and hALOs, which was contrary to the restricted expression pattern of ACE2 (Fig. S2). Therefore, compared to TMPRSS2, ACE2 was the major factor that determined the cell tropism of SARS-CoV-2 in human lung organoids.

Next, we checked whether SARS-CoV-2 infection was associated with proliferation status by co-immunostaining with viral N protein and Ki67 (cycling marker). We found that infected cells (NP^+^) contained both cycling (Ki67^+^) and noncycling (Ki67^-^) cells in hAWOs and most infected cells were cycling cells in hALOs (Fig. S3a). We then checked whether SARS-CoV-2 infection induced apoptosis by co-immunostaining with viral N protein and cleaved Caspase3 (C-Caspas3, apoptotic cell marker). No obvious cell death was observed at 24 hpi or 48 hpi, but at 72 hpi, apoptosis became prominent in both organoids, particularly more in hALOs (Fig. S3b-d).

### Characteristics of SARS-CoV-2 replication in human lung organoids

To conform the viral replication, the ultrastructures of infected hAWOs and hALOs were analyzed by transmission electron microscopy at 72 hpi or 96 hpi. Part of hAWOs and hALOs in one mesh of the grids were shown in Fig. 4a and 4e, and viral particles were found in cells of both organoids (Fig. 4b-d, f and g). In both organoids, viral particles were observed in the apical, lateral and basolateral side of the cells (Fig. 4h-j), indicating potential dissemination route how SARS-CoV-2 passes across the lung epithelial barrier. Double membrane vesicles (DMVs) and convoluted membranes (CMs) with spherules are typical coronavirus replication organelles^42,43^, which were observed in the lung organoids (Fig. 4k). Virus particles in cells were seen in membrane bound vesicles, either as single particles or as groups in enlarged vesicles (Fig. 4l). Enveloped viruses were observed in the lumen of Golgi apparatus and secretory vesicles (Fig. 4m, n), which was consistent with previous report that coronaviruses assembled and matured at the endoplasmic reticulum-Golgi intermediate compartment (ERGIC) and the mature virions were transported to the cell surface and released from the host cells via exocytosis^43,44^. Therefore, TEM analyses captured three critical phases of SARS-CoV-2 life cycle: replication, assembly and release.

**Fig.4.**
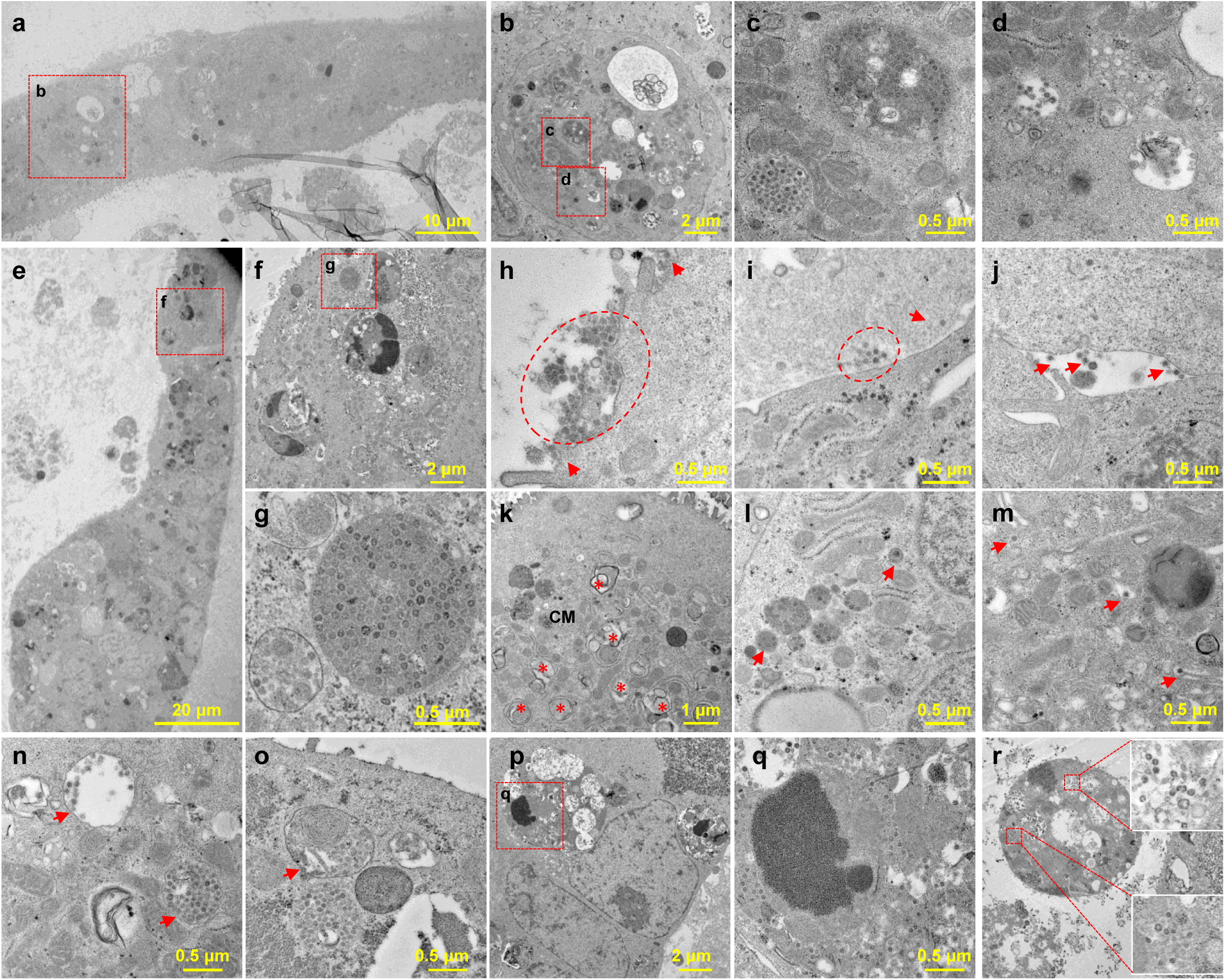
Transmission electron microscopy analysis of SARS-CoV-2 infected human airway and alveolar organoids. **a-d**, Infected hAWOs were fixed and observed under TEM at 96h post infection. A part of the organoids in one mesh was overviewed (**a**) and the virus particles in an infected cell were shown (**b-d**). **e-g**, Infected hALOs were fixed at 72h post infection. A part of the organoids in one mesh (**e**) and the virus particles in an infected cell (**f, g**) were shown. **h-r**, Representative virus particles and typical structures induced by virus infection in hAWOs (**h-n**) and hALOs (**o-r**). Virus particles outside cells at the apical (**h**), basolateral (**i**) and lateral side (**j**). Typical coronavirus replication organelle including double membrane vesicles (DMVs, indicated by asterisks) and convoluted membranes (CMs) with spherules (**k**). Membrane-bound vesicles with one or groups of virus particles (**l**). Enveloped virus particles in Golgi apparatus (**m**). Enveloped virus particles in secretory vesicles (**n**). Virus particles in a lamella body (**o**). Virus particles in a late endosome with engulfed cell debris (**p, q**). Virus particles in disintegrated dead cells (**r**).

Interestingly, we found virus particles within lamellar bodies (Fig. 4o), the typical organelles in AT2 cells, which are essential for pulmonary surfactant synthesis and secretion^45^. Does SARS-CoV-2 hijack lamellar bodies for virus release? Or does SARS-CoV-2 impair the function of lamellar bodies and then the homeostasis of pulmonary surfactant in the alveoli? These questions remain open for further investigation. Additionally, vesicles full of dense virus particles were routinely observed (Fig. 4b, g and n). Besides, virus particles were found in late endosomes with engulfed cell debris (Fig. 4p, q). And more dying cells and engulfed cell debris were observed in hALOs than in hAWOs (Fig. 4r). The TEM data (Fig. 4p-r), as well as the C-Caspase3 immunostaining data (Fig. S3b-d), indicated that the pathological changes of alveoli and bronchioles after SARS-CoV-2 infection were different.

### Early cell response to SARS-CoV-2 infection

To determine the early cell response to SARS-CoV-2 infection, we performed RNA-sequencing analysis using hAWOs and hALOs 48h after SARS-CoV-2 infection. Abundant SARS-CoV-2 viral RNA was detected solely in the infected organoids (Fig. 5a). Principle component analysis (PCA) showed that the samples formed four separate clusters according to organoid type and virus infection (Fig. 5b). In total, 1679 differential expressed genes were identified with 718 genes upregulated and 961 genes downregulated in hAWOs, and 719 genes differential expressed in hALOs with 334 upregulated and 385 downregulated (Fig. 5c). Gene ontology (GO) analysis revealed that most downregulated genes were associated with lipid metabolism, while upregulated genes were associated with immune response (Fig. 5d). Several cytokines and chemokines, including interleukin (IL)-6, tumor necrosis factor (TNF), *CXCL8, CXCL2, CXCL3, CXCL10, CXCL11*, as well as NF-kB related mRNA *NFKB1, NFKB2* and *RELB*, interferon-stimulated genes *ATF3, GEM, IFITM3* and *MX1* were upregulated, consistent with observation in COVID-19 patients^46-48^ (Fig. 5e). ACE2 is the receptor for SARS-CoV and SARS-CoV-2, and SARS-CoV spike (S) protein can induce shedding of ACE2 by ADAM17, which is believed to be a crucial mechanism for SARS-CoV-induced lung injury^49-52^. Surprisingly, we found that the mRNA expression level of *ACE2* was downregulated at 48h after SARS-CoV-2 infection (Fig. 5f). Since most infected cells were viable at 48 hpi (Fig. S3b-d), the downregulation of *ACE2* mRNA was not a secondary effect of cell death but a direct effect of virus infection. Therefore, we believe that SARS-CoV-2 infection might decrease the expression of *ACE2* at both protein and mRNA levels. However, the mechanisms of downregulation remain open for further investigation. In addition, we found that the expression of *TMPRSS2* was also slightly downregulated after SARS-CoV-2 infection at a much less extent than *ACE2* (Fig. 5f).

**Fig.5.**
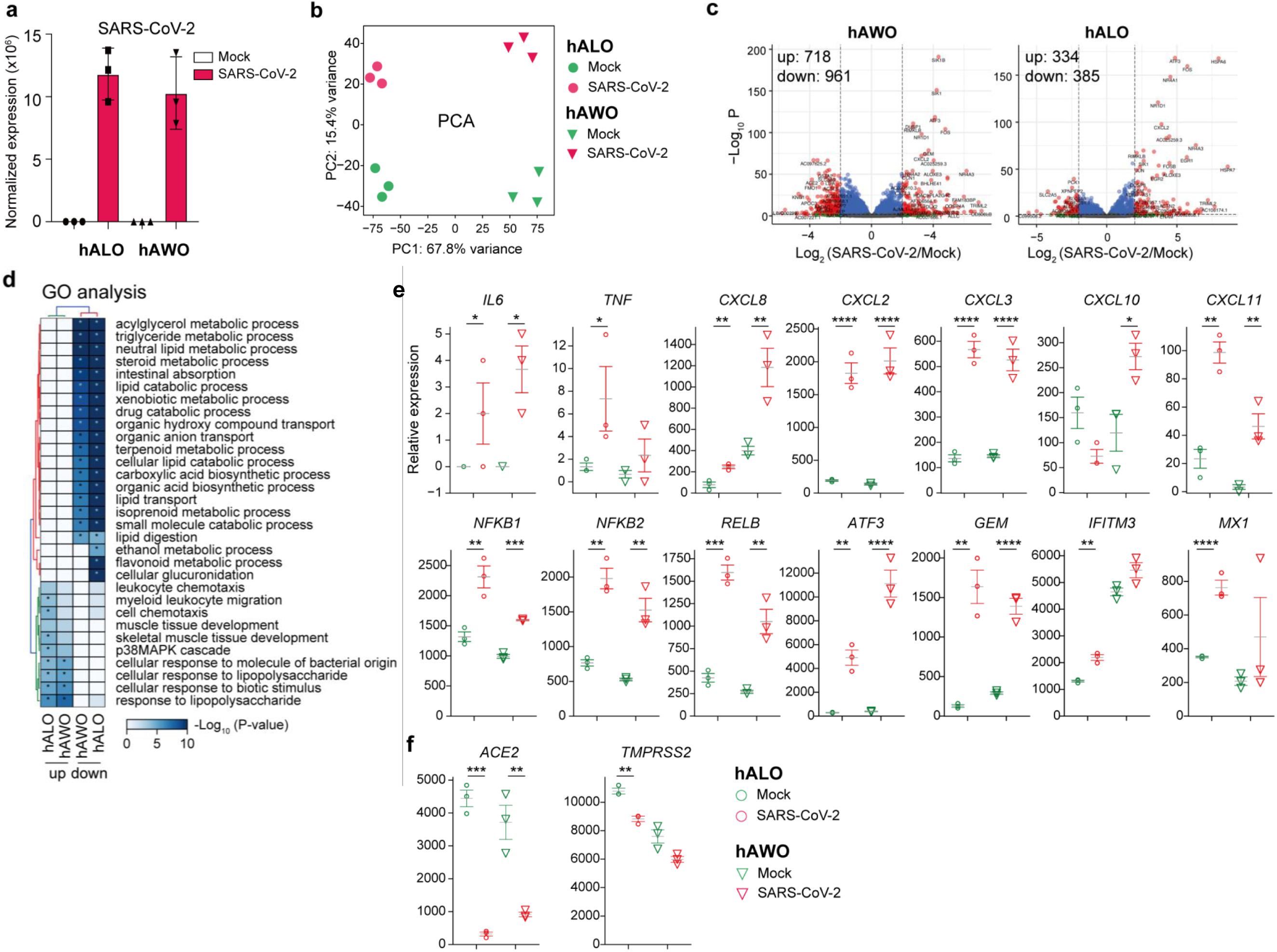
Differentially expressed genes in the SARS-CoV-2-infected human lung organoids. **a**, SARS-CoV-2 viral RNA detected by RNA-seq in mock and infected organoids. Data are expressed as normalized read counts. **b**, PCA plot for the Mock and SARS-CoV-2 infected organoids. **c**, Volcano plot showing differentially expressed genes in the SARS-CoV-2 infected organoids compared with mock control. **d**, Gene ontology (GO) analysis showing the differentially expressed genes from panel c. **e**, Expression level of indicated genes, The grey lines are the means of the three biological replicates, and the error bars are the standard error of the mean. Data are expressed as normalized read counts. P-values are from a one-tailed Student’s t test. * *p*<0.05, ** *p* <0.01, *** *p* <0.001, **** *p* <0.0001. **f**, Same as panel e, but showing the expression of ACE2 and TMPRSS2.

### Remdesivir inhibits SARS-CoV-2 replication inhuman lung organoids

Finally, we tested the inhibitory effect of remdesivir, camostat, and bestatin on the infection of human lung organoids by SARS-CoV-2. Remdesivir is a nucleotide analogue prodrug to inhibit viral replication^53^, which has been reported to repress SARS-CoV-2 infection in basic research and clinic trials^12,54,55^. Camostat is an inhibitor of the serine protease TMPRSS2 that cleaves SARS-CoV-2 S protein and facilitates viral entry^5^. Bestatin is an inhibitor of CD13 (Aminopeptidase N/APN)^56^, a receptor utilized by many α-coronaviruses (SARS-CoV-2 belongs to β-coronaviruses)^44^. As shown in Fig. 6a, remdesivir reduced the production of infectious virus in hAWOs and hALOs, and camostat showed a slightly inhibitory effect in hAWOs not in hALOs, while bestatin had no effects in either hAWOs or hALOs. Quantitative RT-PCR analyses of supernatant viral RNA also demonstrated that remdesivir inhibited viral load (Fig. 6b). We noted that remdesivir reduced viral load to 1/10 but infectious virus titer to less than 1/1000. Similar phenomena, with potent inhibitory effect on virus titer and much less effect on viral load, have been reported in remdesivir treated rhesus macaques with SARS-CoV-2 infection^16^. An explanation for the phenomena might be that virus particles with RNA containing the remdesivir-metabolized adenine analogue are defective for infection, in addition to the known mechanism that remdesivir induces delayed chain termination^53^.

**Fig.6.**
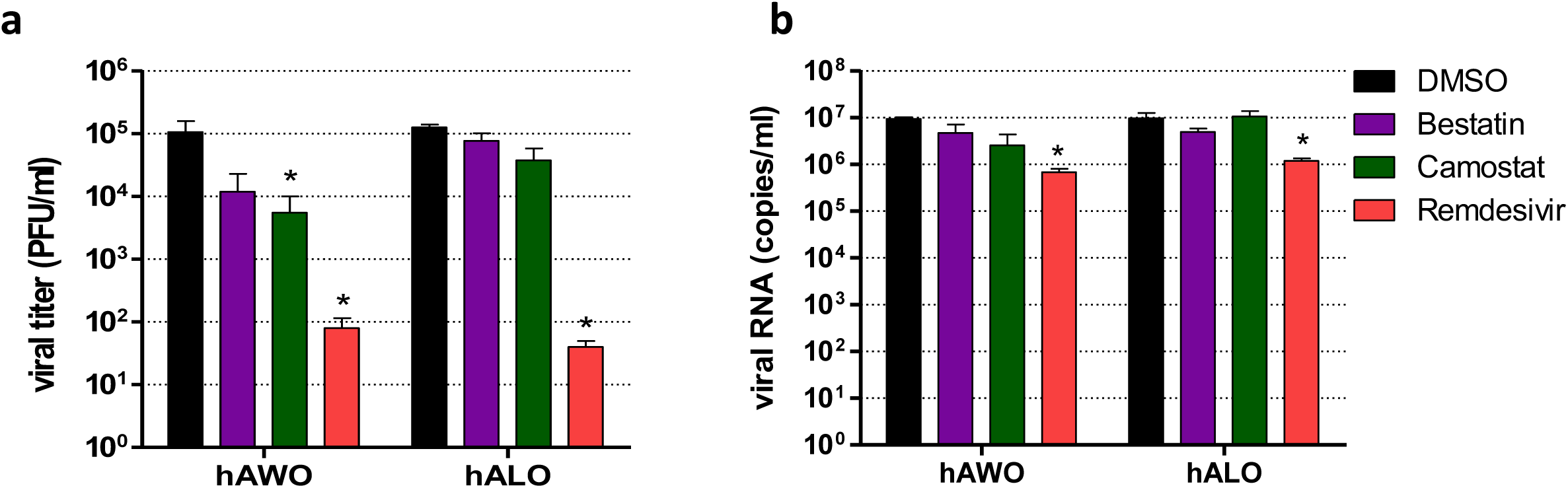
Remdesivir inhibits SARS-CoV-2 replication in both human airway and alveolar organoids. **a, b**, hAWOs and hALOs were infected with SARS-CoV-2, the indicated compounds were added into the culture media 2h post infection. 48h later, the virus titers were determined by plaque assay with Vero E6 cells (**a**) and viral RNA in the culture supernatant was determined by qRT-PCR (**b**).

In summary, we demonstrated that hESCs-derived airway and alveolar organoids could be infected by SARS-CoV-2 and be used for drug test, serving as a pathophysiological model to complement cell lines and animals.

## Acknowledgements

We thank Prof. Mengfeng Li from Southern Medical University for helpful discussion. We are particularly grateful to Tao Du, Lun Wang and the running team from Zhengdian Biosafety Level 3 Laboratory, to Pei Zhang and Anna Du from the core facility of Wuhan Institute of Virology for technical support for TEM experiment. We thank Prof. Zhengli Shi for providing the rabbit antibody against viral N protein. This work was supported by grants from National Natural Science Foundation of China (81872511 and 81670093 to Z.R.), Frontier Research Program of Bioland Laboratory (Guangzhou Regenerative Medicine and Health Guangdong Laboratory) (2018GZR110105005 to Z.R.), National Science and Technology Major Project (2018ZX10301101 to Z.R.), the Natural Science Foundation of Guangdong Province (2018A030313455 to Y.L.), the Program of Department of Science and Technology of Guangdong Province (2014B020212018 to Z.R.), National Key Research and Development Project (2018YFA0507201 to X.C), the special project for COVID-19 of Guangzhou Regenerative Medicine and Health Guangdong Laboratory (2020GZR110106006 to X.C. and J.C.), the emergency grants for prevention and control of SARS-CoV-2 of Guangdong province (2020B111108001 to X.C.) and National Postdoctoral Program for Innovative Talent (BX20190089 to X.Y.)

## Author Contributions

X.C., Z.R., and J.C. initiated, designed and supervised this study; R.P. performed virus infection, viral titer determination, TEM, and drug test experiments; J.F. generated lung organoids and performed immunostaining experiments; X.Y. performed RNA-seq experiment; J.H. analyzed RNA-seq data; Y.Z. and H.S. helped R.P. for virus infection experiments in P3 laboratory; L.L. helped J.F. for immunostaining experiments; S.X. cultured Vero E6 cells; J.X. extracted RNA and performed qRT-PCR experiments; K.W. and H.Z. provided several antibodies for viral N protein; Z.R., J.F., Y.L., R.P., J.C., and X.C. wrote the manuscript.

## Methods

### Maintenance of human ESCs

All experiments in the present study were performed on H9 human embryonic stem cells (hESCs). hESCs were maintained in feeder-free culture conditions in 6-well tissue culture dishes on Matrigel (BD Biosciences, 354277) in mTeSR1 medium (Stem Cell Technologies, 05850) at 37°C with 5% CO_2_. Cells were passaged with TrypLE (Gibco) at 1:6 to 1:8 split ratios every 4 days.

### Generation of hESCs derived hAWO and hALO

hESCs derived hAWOs and hALOs were generated as previously described with modifications^1-3^. H9 cells (∼90% confluence) were cultured in 24-well tissue dishes for 3 days in RPMI1640 medium supplemented with 100ng/ml Activin A (R&D Systems, 338-AC-050) and 2µM CHIR99021 (Tocris, 4423-10MG), followed by 4 days with 200ng/ml Noggin (R&D Systems, 6057-NG-100), 500ng/ml FGF4 (Peprotech, 100-31-1MG), 2µM CHIR99021 and 10µM SB431542 (Tocris, 1614-10MG) in Advanced DMEM/F12 (Life Technologies, 12634010). After 7 days treatment with above-mentioned factors, anterior foregut endodermal cells were embedded in a droplet of Matrigel (BD Biosciences, 356237) and incubated at 37°C with 5% CO_2_ for 20-25 min. After matrigel solidification, cells were then fed with 20ng/ml human BMP4 (R&D Systems, PRD314-10), 0.5µM all-trans retinoic acid (ATRA, Sigma-Aldrich, R2625), 3.5µM CHIR in DMEM/F12 (Life Technologies, 11320033) with 1% Glutamax (Gibco, 35050061), 2% B27 supplement (Life Technologies, 17504044) basal medium from day 8 to day 14. For preconditioning toward lung progenitor stem cell differentiation, NKX2-1^+^ VAFE-enriched cells were cultured in the same basal medium supplemented with 3µM CHIR99021, 10ng/ml human FGF10 (R&D Systems, 345-FG-025), 10ng/ml human KGF (novoprotein, CM88) and 20 µM DAPT (Sigma, D5942) from day 14 to day 21. From day21, human airway organoids (hAWOs) medium was prepared from Ham’s F12 (Gibco, 21127022) by supplementation with 50 nM dexamethasone (Sigma-Aldrich, D4902), 100 nM 8-Br-cAMP (Biolog Life Science Institute, B007-500), 100 nM 3-isobutyl-1-methylxanthine (Wako, 095-03413), 10 ng/ml KGF, 1% B-27 supplement, 0.25% BSA (Sigma, A1470) and 0.1% ITS premix (Corning, 354351). And human alveolar organoids (hALOs) medium was prepared by supplementing 3µM CHIR99021 and 10µM SB431542 to the human airway organoids medium. Organoids were transferred into new Matrigel droplets every 4-7 days using mechanical digestion.

### Quantitative RT-PCR

Total RNA was extracted using the Trizol reagent (MRC, TR1187) and cDNA was converted from 1μg total RNA using the ReverTraAce Kit (TOYOBO, 34520B1). The qPCR reactions were done on Roche LightCycler® 96 PCR system with the SYBR Premix Ex Taq™ Kit (TAKARA, RR420A). Gene expression levels were normalized to *GAPDH* and compared to gene expression levels in hESCs. Three or more biological replicates were performed for each assay and data bars represent mean ± SD. Primers used in this study are listed in Supplementary Table S1.

### SARS-CoV-2 Infection, drug test, and virus titers determination

SARS-CoV-2 (WIV04)^4^ was propagated 7 times on Vero E6 cells in DMEM (Gibico, C12430500BT) with 2% FBS (Gibico, 10099-141) at 37°C with 5% CO_2_. The SARS-CoV-2 isolate was obtained and titrated by plaque assay on Vero E6 cells. Human airway and alveolar organoids were harvested, sheared and resuspended in Ham’s F12 medium (Gibco, 21127022) and infected with virus at multiplicity of infection (MOI) of 1. After 2 hours of SARS-CoV-2 virus adsorption at 37°C in the incubator, cultures were washed twice with Ham’s F12 medium to remove unbound viruses. hAWOs and hALOs were re-embedded into Matrigel (BD Biosciences, 356237) in 24-well tissue plates, and cultured in 500 μL corresponding organoid media, respectively. In drug testing experiments, different drugs at concentration of 10µM were added to the culture 2h after virus infection. Samples were harvested at indicated time points by collecting the supernatant in the wells and the cells via resuspending the matrigel droplet containing organoids into 500 μL Ham’s F12 medium. The viral RNA in the supernatants was extracted by Magnetic Beads Virus RNA Extraction Kit (Shanghai Finegene Biotech, FG438). The intracellular RNA was extracted with Trizol reagent (Invitrogen, 15596026). The viral RNA was quantified by real-time qPCR with Taqman probe targeting the RBD region of S gene. Viral titers (TCID50 equivalants per mL) were determined by plaque assay on Vero E6 cells.

### RNA-seq sequencing and data analysis

Total RNA in the cells was extracted using Trizol (Invitrogen, 15596026) according to the manufacturer’s protocol, and 1ug RNA was used to reverse transcribed into cDNA using Oligo (dT). Fragmented RNA (average length approximately 200 bp) was subjected to first strand and second strand cDNA synthesis followed by adaptor ligation and enrichment with a low-cycle according to the instructions of NEBNext” UltraTM RNA Library Prep Kit for Illumina (NEB, USA). The purified library products were evaluated using the Agilent 2200 TapeStation and Qubit”2.0 (Life Technologies, USA).

Reads were aligned to the human reference genome hg38 with bowtie2^5^, and RSEM^6^ was used to quantify the reads mapped to each gene. Gene expression was normalized by EDASEQ^7^. Differentially expressed genes were obtained using DESeq2 (version 1.10.1)^8^, a cutoff of Q-value < 0.05 and log2 (fold-change) > 1 was used for identify differentially expressed genes. All differentially expressed mRNAs were selected for GO analyses clusterProfiler^9^. Other analysis was performed using glbase^10^. The RNA-seq supporting this study is available at GEO under GSE155717. Data are accessible with a reviewer token: “mbcxaucmpbwttup”.

### Immunofluorescence Staining

For immunofluorescence staining, samples were transferred into 1.5ml tubes and fixed with 4% paraformaldehyde overnight at 4°C or 2h at RT. Following fixation, paraformaldehyde was removed the organoids were rinsed three times with PBS, then the samples were overlaid with O.C.T compound and frozen in liquid nitrogen. The frozen samples were cryosectioned into 6μm sections, washed with PBS three times and permeabilized with 0.2% Triton X-100 (Sigma, T9284)/PBS for 20 min at RT, rinsed again with PBS and then blocked with 5%BSA at RT for 1 hour. The samples were incubated with primary antibodies overnight at 4°C, and then stained with secondary antibodies at RT for 40min. Nuclear counterstained with DAPI (Sigma, D9542) for 3 min, then covered with glass microscope slides and imaged with the Nikon A1 confocal microscope. NIS-Elements software was used to render Z-stack three-dimensional images. The primary and secondary antibodies used in this study are listed in Supplementary Table S2.

### Transmission Electron Microscopy

Organoids were collected and fixed in 2.5% glutaraldehyde for 24h, washed with 0.1M Phosphate buffer (19ml 0.2M NaH_2_PO_4_, 81ml 0.2 M Na_2_HPO_4_) for 3 times, and further fixed with 1% Osimium tetraoxide for 2h at room temperature. The fixed organoids were then washed with phosphate buffer and dehydrated with 30%, 50%, 70%, 80%, 85%, 90%, 95%, and 100% alcohol sequentially. After a step of infiltration with different mixtures of acetone-epon (2:1, 1:1, vol/vol), the samples were embedded in pure Epon. Polymerization was performed by incubation at 60°C for 48h. Ultra-thin sections (80-100 nm) were cut on Ultramicrotome (Leica EM UC7), put on grids and stained with uranyl acetate and lead citrate. After wash and drying, images were acquired by the digital camera on TEM (FEI, Tecnai G2 20 TWIN, 200kv), with identical magnificence.

### Experimental replicates and statistical analysis

Error bars in these figures indicate S.D. (for qRT-PCR) and S.E.M (for other assays) Unpaired, two-tailed Student’s t tests were used for comparisons between two groups of n=3 or more samples. P<0.05 was defined as statistical significance. Immunofluorescence (IF) imaging were done on Z-stacks acquired with confocal microscope at least three (n=3) independent biological samples or more. The co-localization of quantitative analysis of specific immunofluorescence marker was shown in figure legends. All of the statistical analyses in this study were done with GraphPad Prism 8 software.

---

**Extended Data Fig.1.**
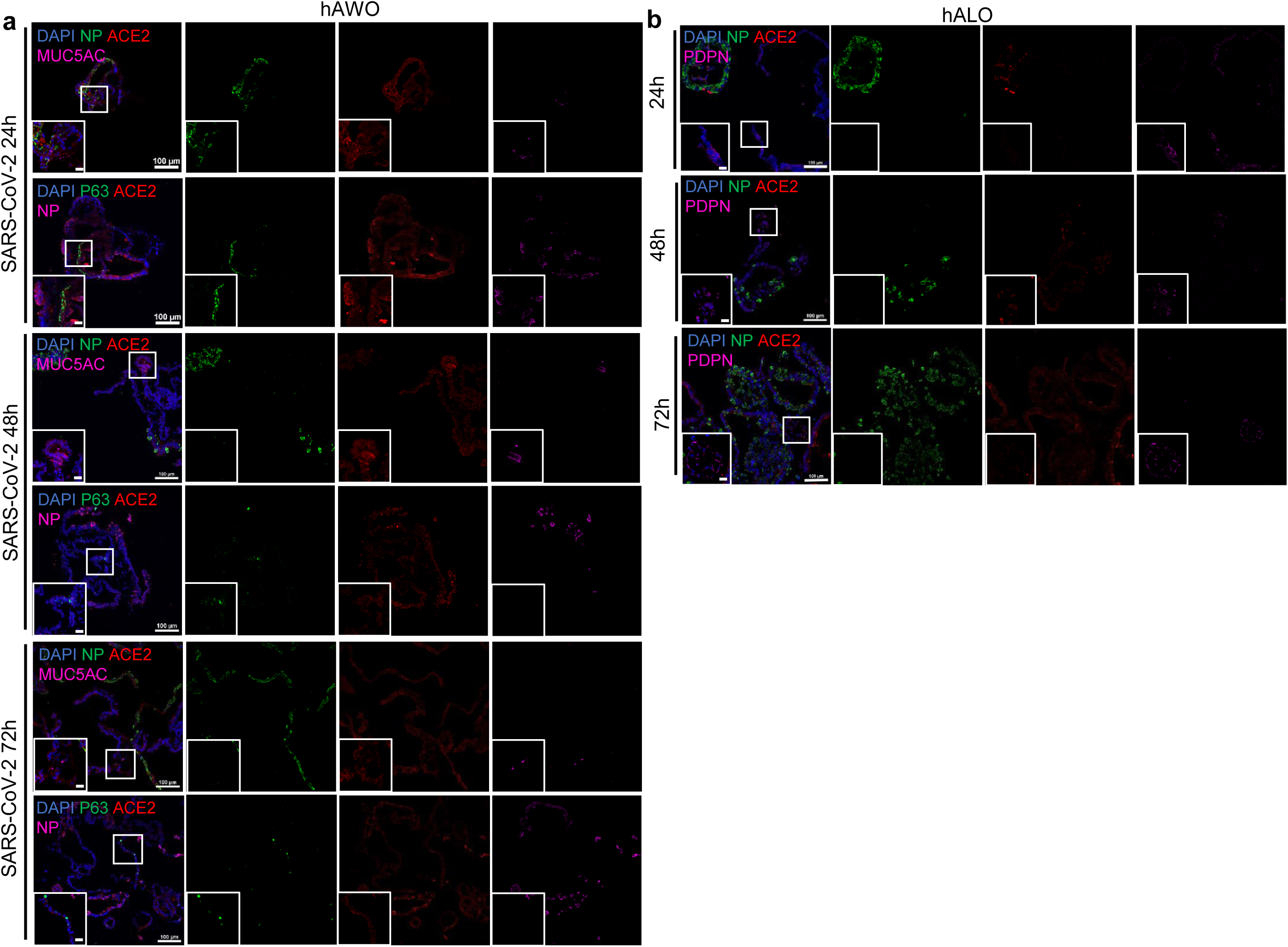
SARS-CoV-2 dose not infect basal cells, goblet cells or alveolar type I cells. **a**,**b**, Representative immunofluorescence images of nucleoprotein, ACE2 and indicated cell linage marker expression with DNA stain (DAPI). Basal cells (P63^+^) and goblet cells (MUC5AC^+^) were stained in human airway organoids at indicated time points (**a**). Alveolar type I cells (PDPN^+^) were stained in human alveolar organoids (**b**). Scale bar, 100µm; bottom left corner, 20µm. Boxes represent zoom views.

**Extended Data Fig.2.**
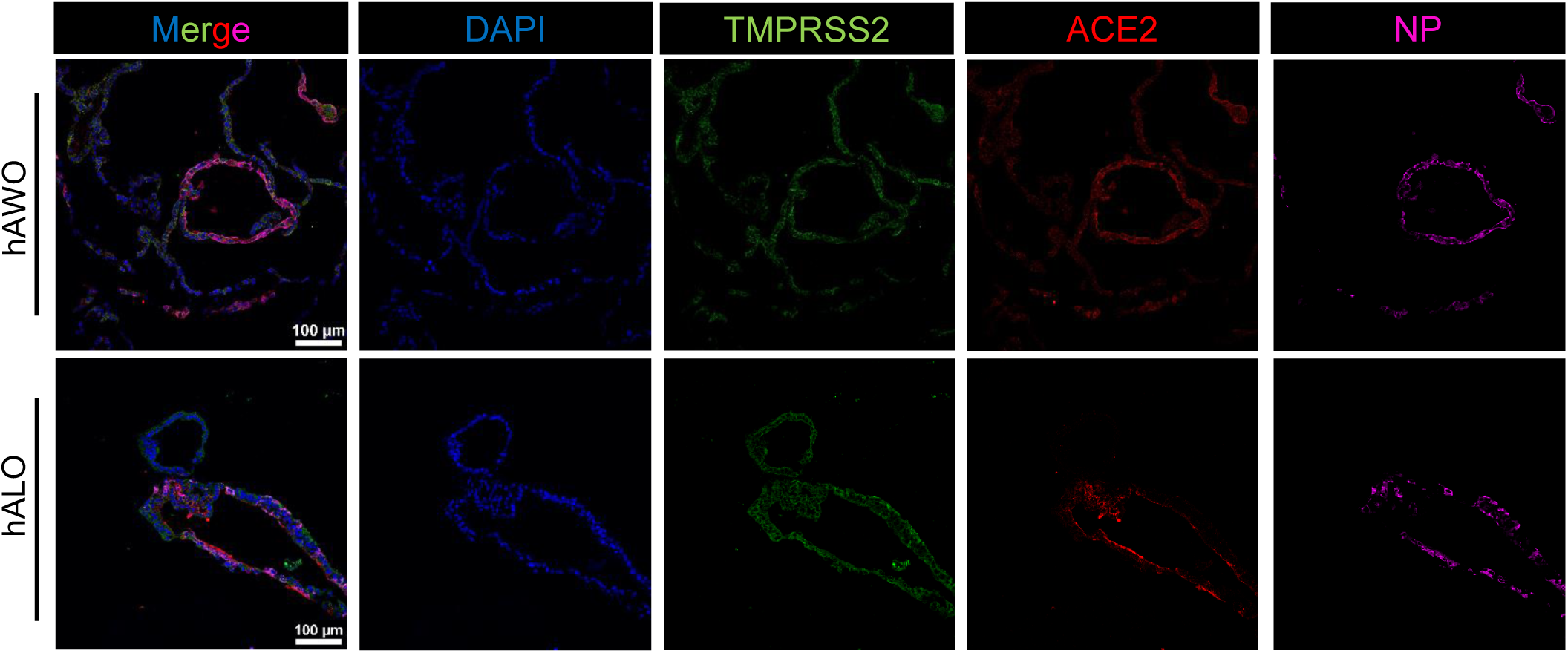
TMPRSS2 is ubiquitously expressed in human airway and alveolar organoid cells. Immunofluorescence images of SARS-CoV-2 infected human airway and alveolar organoids. TMPRSS2 (green) is broadly expressed in almost all human lung epithelial cells. Virus infected cells (nucleoprotein positively) highly express ACE2 (red). Scale bar, 100µm.

**Extended Data Fig.3.**
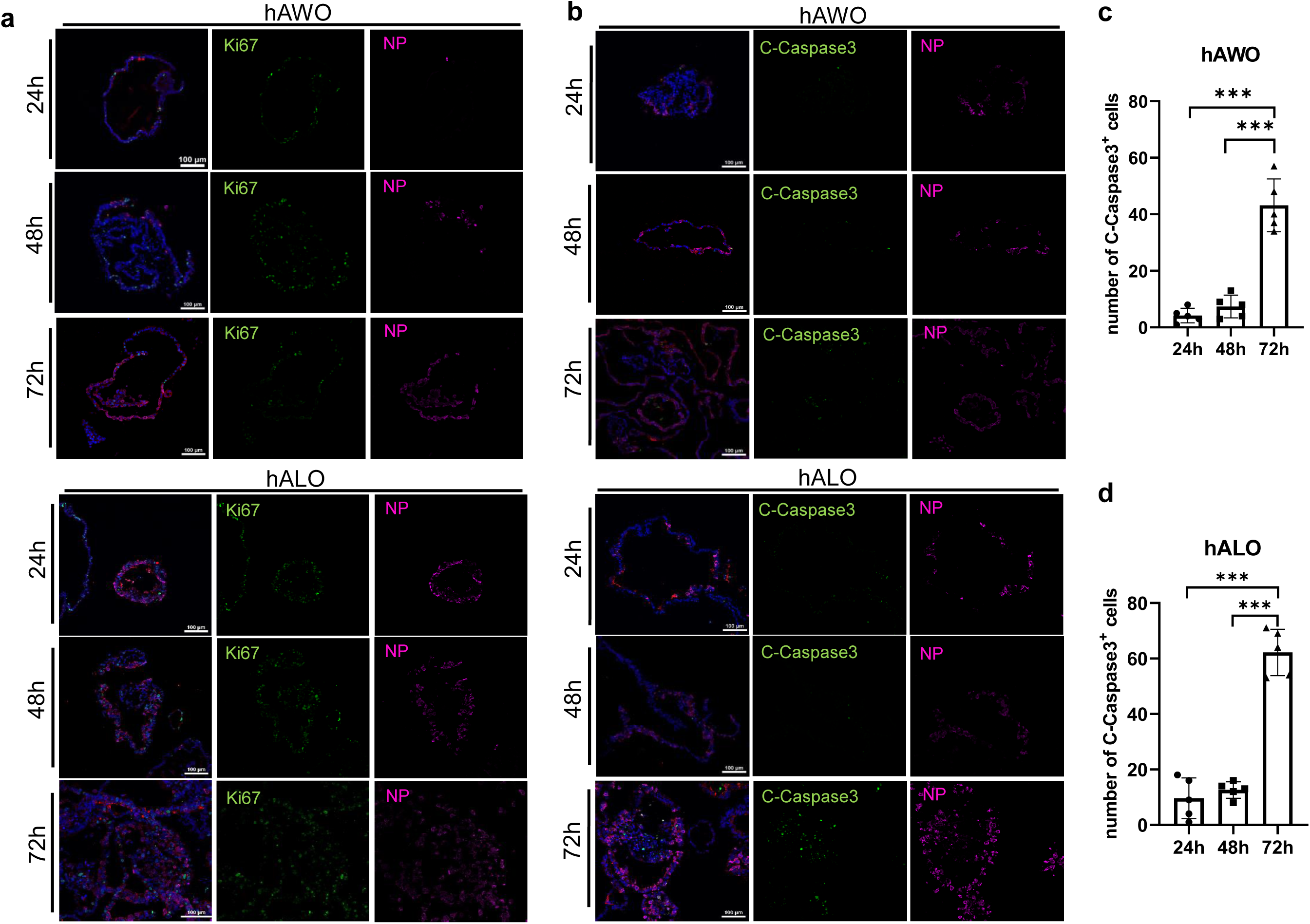
SRAS-CoV-2 infection induces apoptosis in human airway and alveolar organoids. **a**, SARS-CoV-2 infected human airway and alveolar organoids are stained by cell proliferation marker, Ki67 (green) at 24, 48 and 72 hpi. Scale bar, 100µm. **b**, Long term infection of SARS-CoV-2 induces apoptosis. Cleaved caspase-3 (green) were observed within virus infected organoids at 72 hpi. Scale bar, 100µm. **c**,**d**, Number of cleaved caspase-3 positive cells in SARS-CoV-2 infected human airway organoids(**c**) and human alveolar organoids(**d**). n=5 organoids per condition. *** *p* <0.001 by unpaired, two-tailed Student’s t test.

**Table S1.**
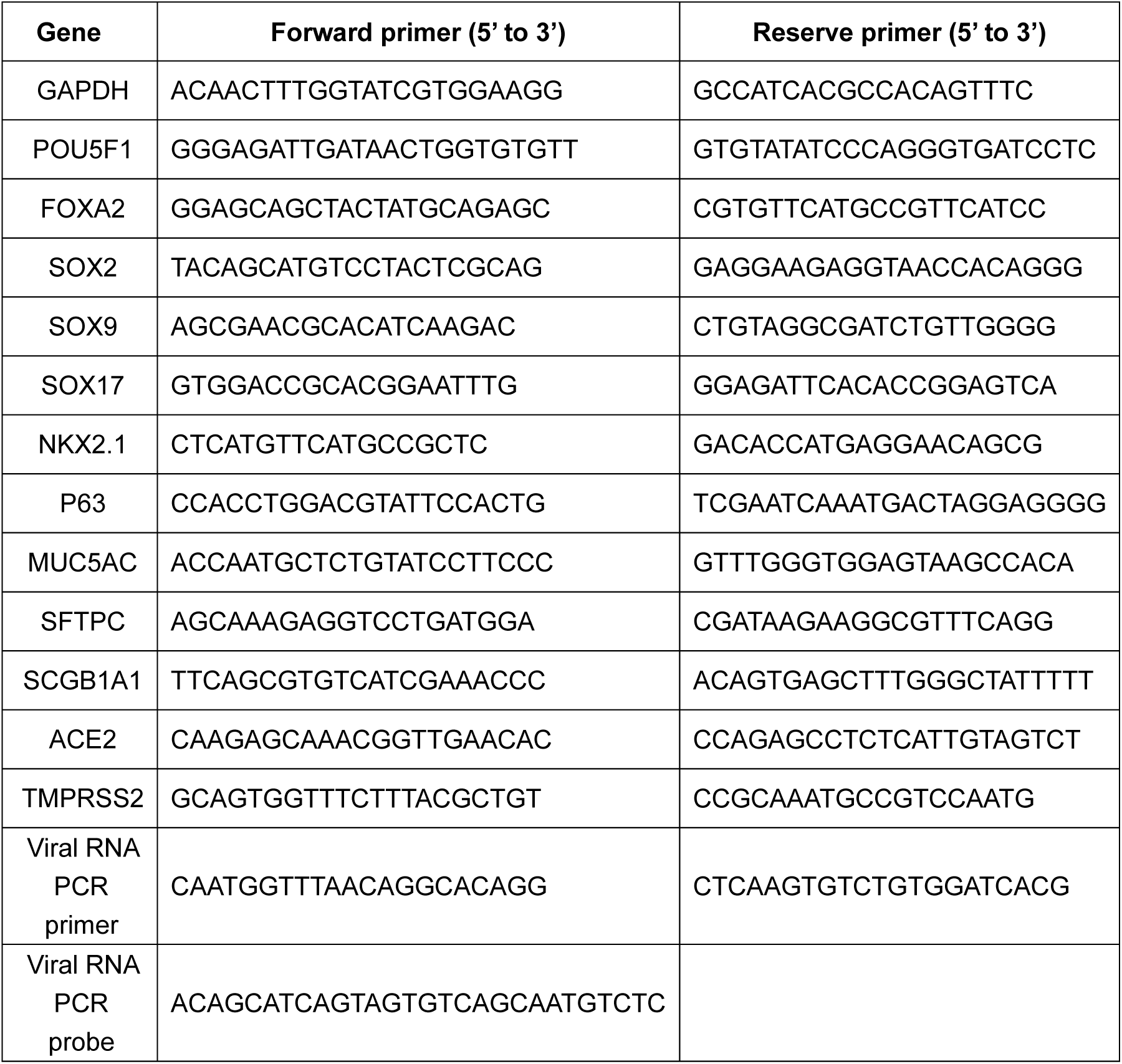
Primers for qRT-PCR.

**Table S2.** Antibody list.

| Primary Antibodies | Dilution rate | Manufacturer | Cat. No. |
| --- | --- | --- | --- |
| NKX2.1 | 1:250 | Abcam | ab76013 |
| SOX2 | 1:1000 | Abcam | AB97959 |
| SOX9 | 1:40 | R&D systems | AF3075 |
| P63 | 1:200 | Abcam | ab124762 |
| MUC5AC | 1:150 | Thermo Fisher Scientific | MA5-12178 |
| CC10 | 1:300 | Abcam | Ab40873 |
| SFTPC | 1:300 | SEVEN HILLS | WRAB-76694 |
| AQP5 | 1:150 | Abcam | ab92320 |
| PDPN | 1:200 | Abcam | ab10288 |
| acetylated Tubulin | 1:1000 | Sigma | T7451 |
| Pro-SPC | 1:200 | EMD-Millipore | #AB3786 |
| E-CAD | 1:100 | R&D systems | AF748 |
| Ki67 | 1:250 | Abcam | Ab1667 |
| Cleaved Caspase-3 | 1:400 | Cell Signaling Technology | #9661 |
| Human ACE-2 | 1:100 | R&D systems | AF933 |
| TMPRSS2 | 1:150 | Abcam | Ab109131 |
| SARS-CoV-2<br>Nucleocapsid | 1:200 | Sino biological | 40143-MM08 |
| SARS-CoV-2<br>Nucleocapsid | 1:5000 | Kindly provide by Prof.<br>Zheng-Li Shi | Reference<sup>1</sup> |
| Dnokey anti-goat<br>( RRX ) | 1:500 | Jackson ImmunoResearch | 705-295-147 |
| Donkey anti-rabbit<br>(Alexa488) | 1:500 | Thermo Fisher Scientific | A-21206 |
| Donkey anti-mouse<br>(Alexa647) | 1:300 | Thermo Fisher Scientific | A-31571 |

## References

1 Li, J. Y. et al. The epidemic of 2019-novel-coronavirus (2019-nCoV) pneumonia and insights for emerging infectious diseases in the future. Microbes Infect, doi: 10.1016/j.micinf.2020.02.002 (2020).

2 Wiersinga, W. J., Rhodes, A., Cheng, A. C., Peacock, S. J. & Prescott, H. C. Pathophysiology, Transmission, Diagnosis, and Treatment of Coronavirus Disease 2019 (COVID-19): A Review. JAMA, doi: 10.1001/jama.2020.12839 (2020).

3 Zhu, N. et al. A Novel Coronavirus from Patients with Pneumonia in China, 2019. N Engl J Med 382, 727–733, doi: 10.1056/NEJMoa2001017 (2020).

4 Takayama, K. In Vitro and Animal Models for SARS-CoV-2 research. Trends Pharmacol Sci 41, 513–517, doi: 10.1016/j.tips.2020.05.005 (2020).

5 Hoffmann, M. et al. SARS-CoV-2 Cell Entry Depends on ACE2 and TMPRSS2 and Is Blocked by a Clinically Proven Protease Inhibitor. Cell 181, 271–280 e278, doi: 10.1016/j.cell.2020.02.052 (2020).

6 Kaye, M. et al. SARS-associated coronavirus replication in cell lines. Emerg Infect Dis 12, 128–133, doi:DOI 10.3201/eid1201.050496 (2006).

7 Song, Z. et al. From SARS to MERS, Thrusting Coronaviruses into the Spotlight. Viruses 11, doi: 10.3390/v11010059 (2019).

8 Harcourt, J. et al. Isolation and characterization of SARS-CoV-2 from the first US COVID-19 patient. bioRxiv, doi: 10.1101/2020.03.02.972935 (2020).

9 Zhou, P. et al. A pneumonia outbreak associated with a new coronavirus of probable bat origin. Nature 579, 270–273, doi: 10.1038/s41586-020-2012-7 (2020).

10 Kim, J. M. et al. Identification of Coronavirus Isolated from a Patient in Korea with COVID-19. Osong Public Health Res Perspect 11, 3–7, doi: 10.24171/j.phrp.2020.11.1.02 (2020).

11 Ou, X. et al. Characterization of spike glycoprotein of SARS-CoV-2 on virus entry and its immune cross-reactivity with SARS-CoV. Nat Commun 11, 1620, doi: 10.1038/s41467-020-15562-9 (2020).

12 Wang, M. et al. Remdesivir and chloroquine effectively inhibit the recently emerged novel coronavirus (2019-nCoV) in vitro. Cell Res, doi: 10.1038/s41422-020-0282-0 (2020).

13 Bao, L. et al. The pathogenicity of SARS-CoV-2 in hACE2 transgenic mice. Nature, doi: 10.1038/s41586-020-2312-y (2020).

14 Jiang, R. D. et al. Pathogenesis of SARS-CoV-2 in Transgenic Mice Expressing Human Angiotensin-Converting Enzyme 2. Cell 182, 50–58 e58, doi: 10.1016/j.cell.2020.05.027 (2020).

15 van Doremalen, N. et al. ChAdOx1 nCoV-19 vaccination prevents SARS-CoV-2 pneumonia in rhesus macaques. bioRxiv, doi: 10.1101/2020.05.13.093195 (2020).

16 Williamson, B. N. et al. Clinical benefit of remdesivir in rhesus macaques infected with SARS-CoV-2. Nature, doi: 10.1038/s41586-020-2423-5 (2020).

17 Chandrashekar, A. et al. SARS-CoV-2 infection protects against rechallenge in rhesus macaques. Science, doi: 10.1126/science.abc4776 (2020).

18 Rockx, B. et al. Comparative pathogenesis of COVID-19, MERS, and SARS in a nonhuman primate model. Science 368, 1012-+, doi: 10.1126/science.abb7314 (2020).

19 Shi, J. Z. et al. Susceptibility of ferrets, cats, dogs, and other domesticated animals to SARS-coronavirus 2. Science 368, 1016-+, doi: 10.1126/science.abb7015 (2020).

20 Yu, J. et al. DNA vaccine protection against SARS-CoV-2 in rhesus macaques. Science, doi: 10.1126/science.abc6284 (2020).

21 Sia, S. F. et al. Pathogenesis and transmission of SARS-CoV-2 in golden hamsters. Nature, doi: 10.1038/s41586-020-2342-5 (2020).

22 Warren, H. S. et al. Mice are not men. Proc Natl Acad Sci U S A 112, E345, doi: 10.1073/pnas.1414857111 (2015).

23 Martic-Kehl, M. I., Schibli, R. & Schubiger, P. A. Can animal data predict human outcome? Problems and pitfalls of translational animal research. Eur J Nucl Med Mol I 39, 1492–1496, doi: 10.1007/s00259-012-2175-z (2012).

24 Sun, D. et al. Comparison of human duodenum and Caco-2 gene expression profiles for 12,000 gene sequences tags and correlation with permeability of 26 drugs. Pharm Res 19, 1400–1416, doi: 10.1023/a:1020483911355 (2002).

25 Pan, C., Kumar, C., Bohl, S., Klingmueller, U. & Mann, M. Comparative proteomic phenotyping of cell lines and primary cells to assess preservation of cell type-specific functions. Mol Cell Proteomics 8, 443–450, doi: 10.1074/mcp.M800258-MCP200 (2009).

26 Cairns, R. A., Harris, I. S. & Mak, T. W. Regulation of cancer cell metabolism. Nat Rev Cancer 11, 85–95, doi: 10.1038/nrc2981 (2011).

27 Clevers, H. Modeling Development and Disease with Organoids. Cell 165, 1586–1597, doi: 10.1016/j.cell.2016.05.082 (2016).

28 Rossi, G., Manfrin, A. & Lutolf, M. P. Progress and potential in organoid research. Nat Rev Genet 19, 671–687, doi: 10.1038/s41576-018-0051-9 (2018).

29 Dutta, D. & Clevers, H. Organoid culture systems to study host-pathogen interactions. Curr Opin Immunol 48, 15–22, doi: 10.1016/j.coi.2017.07.012 (2017).

30 Ramani, S., Crawford, S. E., Blutt, S. E. & Estes, M. K. Human organoid cultures: transformative new tools for human virus studies. Curr Opin Virol 29, 79–86, doi: 10.1016/j.coviro.2018.04.001 (2018).

31 Han, Y. et al. Identification of Candidate COVID-19 Therapeutics using hPSC-derived Lung Organoids. bioRxiv, doi: 10.1101/2020.05.05.079095 (2020).

32 Suzuki, T. et al. Generation of human bronchial organoids for SARS-CoV-2 research. bioRxiv (2020).

33 Monteil, V. et al. Inhibition of SARS-CoV-2 Infections in Engineered Human Tissues Using Clinical-Grade Soluble Human ACE2. Cell 181, 905–913 e907, doi: 10.1016/j.cell.2020.04.004 (2020).

34 Zhou, J. et al. Infection of bat and human intestinal organoids by SARS-CoV-2. Nat Med 26, 1077–1083, doi: 10.1038/s41591-020-0912-6 (2020).

35 Zhao, B. et al. Recapitulation of SARS-CoV-2 infection and cholangiocyte damage with human liver ductal organoids. Protein Cell, doi: 10.1007/s13238-020-00718-6 (2020).

36 Lamers, M. M. et al. SARS-CoV-2 productively infects human gut enterocytes. Science 369, 50–54, doi: 10.1126/science.abc1669 (2020).

37 Yang, L. et al. A Human Pluripotent Stem Cell-based Platform to Study SARS-CoV-2 Tropism and Model Virus Infection in Human Cells and Organoids. Cell Stem Cell 27, 125–136 e127, doi: 10.1016/j.stem.2020.06.015 (2020).

38 Chen, Y. et al. Long-Term Engraftment Promotes Differentiation of Alveolar Epithelial Cells from Human Embryonic Stem Cell Derived Lung Organoids. Stem Cells Dev 27, 1339–1349, doi: 10.1089/scd.2018.0042 (2018).

39 McCauley, K. B. et al. Efficient Derivation of Functional Human Airway Epithelium from Pluripotent Stem Cells via Temporal Regulation of Wnt Signaling. Cell Stem Cell 20, 844–857 e846, doi: 10.1016/j.stem.2017.03.001 (2017).

40 Yamamoto, Y. et al. Long-term expansion of alveolar stem cells derived from human iPS cells in organoids. Nat Methods 14, 1097–1106, doi: 10.1038/nmeth.4448 (2017).

41 Hou, Y. J. et al. SARS-CoV-2 Reverse Genetics Reveals a Variable Infection Gradient in the Respiratory Tract. Cell 182, 429–446 e414, doi: 10.1016/j.cell.2020.05.042 (2020).

42 van Hemert, M. J. et al. SARS-coronavirus replication/transcription complexes are membrane-protected and need a host factor for activity in vitro. PLoS Pathog 4, e1000054, doi: 10.1371/journal.ppat.1000054 (2008).

43 Hilgenfeld, R. & Peiris, M. From SARS to MERS: 10 years of research on highly pathogenic human coronaviruses. Antiviral Res 100, 286–295, doi: 10.1016/j.antiviral.2013.08.015 (2013).

44 Fehr, A. R. & Perlman, S. Coronaviruses: an overview of their replication and pathogenesis. Methods Mol Biol 1282, 1–23, doi: 10.1007/978-1-4939-2438-7_1 (2015).

45 Schmitz, G. & Muller, G. Structure and function of lamellar bodies, lipid-protein complexes involved in storage and secretion of cellular lipids. J Lipid Res 32, 1539–1570 (1991).

46 Huang, C. et al. Clinical features of patients infected with 2019 novel coronavirus in Wuhan, China. The Lancet 395, 497–506, doi: 10.1016/s0140-6736(20)30183-5 (2020).

47 He, J. et al. Single-cell analysis reveals bronchoalveolar epithelial dysfunction in COVID-19 patients. Protein Cell, doi: 10.1007/s13238-020-00752-4 (2020).

48 Wilk, A. J. et al. A single-cell atlas of the peripheral immune response in patients with severe COVID-19. Nature Medicine 26, 1070–1076, doi: 10.1038/s41591-020-0944-y (2020).

49 Glowacka, I. et al. Differential downregulation of ACE2 by the spike proteins of severe acute respiratory syndrome coronavirus and human coronavirus NL63. J Virol 84, 1198–1205, doi: 10.1128/JVI.01248-09 (2010).

50 Vaduganathan, M. et al. Renin-Angiotensin-Aldosterone System Inhibitors in Patients with Covid-19. N Engl J Med 382, 1653–1659, doi: 10.1056/NEJMsr2005760 (2020).

51 Kuba, K. et al. A crucial role of angiotensin converting enzyme 2 (ACE2) in SARS coronavirus-induced lung injury. Nat Med 11, 875–879, doi: 10.1038/nm1267 (2005).

52 Heurich, A. et al. TMPRSS2 and ADAM17 cleave ACE2 differentially and only proteolysis by TMPRSS2 augments entry driven by the severe acute respiratory syndrome coronavirus spike protein. J Virol 88, 1293–1307, doi: 10.1128/JVI.02202-13 (2014).

53 Eastman, R. T. et al. Remdesivir: A Review of Its Discovery and Development Leading to Emergency Use Authorization for Treatment of COVID-19. ACS Cent Sci 6, 672–683, doi: 10.1021/acscentsci.0c00489 (2020).

54 Wang, Y. et al. Remdesivir in adults with severe COVID-19: a randomised, double-blind, placebo-controlled, multicentre trial. Lancet 395, 1569–1578, doi: 10.1016/S0140-6736(20)31022-9 (2020).

55 Beigel, J. H. et al. Remdesivir for the Treatment of Covid-19 - Preliminary Report. N Engl J Med, doi: 10.1056/NEJMoa2007764 (2020).

56 Jia, M. R., Wei, T. & Xu, W. F. The Analgesic Activity of Bestatin as a Potent APN Inhibitor. Front Neurosci 4, 50, doi: 10.3389/fnins.2010.00050 (2010).

## Reference

1 Yamamoto, Y. et al. Long-term expansion of alveolar stem cells derived from human iPS cells in organoids. Nat Methods 14, 1097–1106, doi: 10.1038/nmeth.4448 (2017).

2 McCauley, K. B. et al. Efficient Derivation of Functional Human Airway Epithelium from Pluripotent Stem Cells via Temporal Regulation of Wnt Signaling. Cell Stem Cell 20, 844–857 e846, doi: 10.1016/j.stem.2017.03.001 (2017).

3 Chen, Y. et al. Long-Term Engraftment Promotes Differentiation of Alveolar Epithelial Cells from Human Embryonic Stem Cell Derived Lung Organoids. Stem Cells Dev 27, 1339–1349, doi: 10.1089/scd.2018.0042 (2018).

4 Zhou, P. et al. A pneumonia outbreak associated with a new coronavirus of probable bat origin. Nature 579, 270–273, doi: 10.1038/s41586-020-2012-7 (2020).

5 Langmead, B. & Salzberg, S. L. Fast gapped-read alignment with Bowtie 2. Nat Methods 9, 357–359, doi: 10.1038/nmeth.1923 (2012).

6 Li, B. & Dewey, C. N. RSEM: accurate transcript quantification from RNA-Seq data with or without a reference genome. BMC bioinformatics 12, 323–323, doi: 10.1186/1471-2105-12-323 (2011).

7 Risso, D., Schwartz, K., Sherlock, G. & Dudoit, S. GC-content normalization for RNA-Seq data. BMC bioinformatics 12, 480–480, doi: 10.1186/1471-2105-12-480 (2011).

8 Love, M. I., Huber, W. & Anders, S. Moderated estimation of fold change and dispersion for RNA-seq data with DESeq2. Genome Biol 15, 550, doi: 10.1186/s13059-014-0550-8 (2014).

9 Yu, G., Wang, L.-G., Han, Y. & He, Q.-Y. clusterProfiler: an R package for comparing biological themes among gene clusters. OMICS 16, 284–287, doi: 10.1089/omi.2011.0118 (2012).

10 Hutchins, A. P., Jauch, R., Dyla, M. & Miranda-Saavedra, D. glbase: a framework for combining, analyzing and displaying heterogeneous genomic and high-throughput sequencing data. Cell Regen (Lond) 3, 1, doi: 10.1186/2045-9769-3-1 (2014).

## References

1 Zhou, P. et al. A pneumonia outbreak associated with a new coronavirus of probable bat origin. Nature 579, 270–273, doi: 10.1038/s41586-020-2012-7 (2020).

